# The role of glycaemic and lipid risk factors in mediating the effect of BMI on coronary heart disease: A two-step, two-sample Mendelian randomization study

**DOI:** 10.1101/118133

**Authors:** Lin Xu, Maria Carolina Borges, Gibran Hemani, Deborah A Lawlor

## Abstract

**Background:** The extent to which effects of BMI on coronary heart disease (CHD) are mediated by gylcaemic and lipid risk factors is unclear.

**Methods:** We used two-sample Mendelian randomization to determine the causal effect of: (i) BMI on CHD (60,801 cases; 123, 504 controls), type 2 diabetes (T2DM; 34,840 cases; 114,981 controls), fasting glucose (n=46,186), insulin (n=38,238), HbA_1c_ (n=46,368), LDL-cholesterol (LDL-C), HDL-cholesterol (HDL-C) and triglycerides (n=188,577); (ii) glycaemic and lipids traits on CHD; and (iii) extent to which these traits mediated any effect of BMI on CHD.

**Findings:** One standard deviation (SD) increase in BMI (~ 4.5kg/m^2^) increased CHD (odds ratio=1.45 (95% confidence interval (CI): 1.27, 1.66)) and T2DM (1.96 (1.35, 2.83)), and levels of fasting glucose (0.07mmol/l (95%CI 0.03, 0.11)), HbA_1c_ (0.05% (95%CI 0.01, 0.08)), fasting insulin (0.18log pmol/l (95%CI 0.14, 0.22)) and triglycerides (0.20 SD (95%CI 0.14, 0.26)), and lowered levels of HDL-C (−0.23 SD (95%CI −0.32, −0.15)). BMI was not causally related to LDL-C. After accounting for potential pleiotropy, triglycerides, HbA_1c_ and T2DM were causally related to CHD. The BMI-CHD effect reduced from 1.45 to 1.16 (95%CI 0.99, 1.36) and to 1.36 (95%CI 1.19, 1.57) with genetic adjustment for triglycerides or HbA_1c_ respectively, and to 1.09 (95%CI 0.94, 1.27) with adjustment for both.

**Interpretation:** Increased triglyceride levels and poor glycaemic control appear to mediate much of the effect of BMI on CHD.

**Funding:** European Research Council (669545), European Union (733206), China Medical Board (CMB_2015/16), Conselho Nacional de Desenvolvimento Científico e Tecnológico and UK Medical Research Council (MC_UU_12013/5).

## Research in context

### Evidence before this study

We searched PubMed (https://www.ncbi.nlm.nih.gov/pubmed) using various combinations of the following search terms: (“body mass index” or “BMI” or “adiposity” or “weight”) and (“coronary heart disease” or “CHD” or “myocardial infarction” or “MI” or “cardiovascular disease” or “CVD”) and (“diabetes” or “glucose”or “gylcaeted haemoglobin” or “HbA1c” or “insulin” or “insulin resistance” or “insulin sensitivity”) and (“lipid” or “dyslipidaemia” or “low density lipoprotein cholesterol” or “LDL-C” or “high density lipoprotein cholesterol” or “HDL-C” or “triglyceride” or “cholesterol”) and (“Mendelian” or “Mendelian randomization” or “Mendelian randomisation”) and (“two-step” or “mediation”). We used these searches to identify any published paper (published in English) that was a Mendelian randomization study testing the effect of BMI on CHD. We ran the searches so that they would include any such study that tested mediation (by any factors) of the effect of BMI on CHD, but was not exclusive for mediation studies. We identified four publications, from three studies that fulfilled our criteria. These were all one-sample (both effect of genetic instrumental variable on BMI and of genetic instrumental variable on CHD from the same study sample(s)) and all in predominantly European origin participants. The number of CHD cases in these studies were between 3062 and 11056. Whilst one of these studies reported a null effect of BMI on CHD based on a p-value threshold of 0.05, when we meta-analysed the results there was evidence of a positive effect of BMI from these three studies: Odds ratio of CHD per 4.5kg/m^2^ 1.34 (95%CI: 1.03, 1.74), I^2^ 47%. We identified only one study that used Mendelian randomization to test the effect of BMI on CHD and then explored possible mediators, despite finding no evidence of an effect of BMI on LDL-C they concluded that it explained 8% of the effect of BMI on CHD, and further that remnant cholesterol (triglyceride) explained 7% and systolic blood pressure explained 8%. That study was unable to explore the potential mediating effect of insulin resistance or glycaemic risk factors on the effect of BMI on CHD. This is important as BMI is strongly related to insulin resistance/hyperglycaemia, which in turn is a strong risk factor for CHD.

### Added value of this study

We used 77 independent single nucleotide polymorphisms (SNPs) identified as robustly associated with BMI from genome-wide association studies, as genetic instrumental variables to test the causal effect of BMI on CHD and the extent to which this was mediated by lipid or insulin/glycaemic traits. In comparison with existing evidence, we used 2-sample Mendelian randomisation, were able to explore mediation by insulin/glycaemic traits and explore whether we could replicate the lipid mediating effects from the one previous study to assess this and had considerably larger sample sizes (e.g. 60,801 CHD cases). Importantly, we were able to apply a number of sensitivity analyses using novel methods for testing possible bias due to pleiotopy (the likely key source of bias in Mendelian randomization studies) that have been developed for use in 2-sample Mendelian randomization. We found evidence for a positive effect of BMI on CHD, type 2 diabetes, fasting glucose, HbA_1c_, fasting insulin and triglycerides, and an inverse effect on HDL-C; we did not find robust evidence for an effect of BMI on LDL-C. After accounting for potential pleiotropy, we also found that triglycerides, HbA_1c_ and T2DM were causally related to CHD, and that these mediated most of the effect of BMI on CHD..

### Implications of all the available evidence

Whilst preventing overweight and obesity is an important public health aim, the substantial and increasing number of people with a high body mass index highlights the need for secondary prevention that aims to reduce risk of the main disease outcomes of high BMI (e.g. CHD) in those with high BMI, by targeting causal mediators. Treating causal mediators of the effect of BMI on CHD could mitigate its effect, but biases in conventional epidemiological methods for testing mediation have limited our understanding of which CHD risk factors mediate BMI effects. Our findings provide strong support for undertaking RCTs in obese people to test the effect of triglyceride reduction and glycaemic control as a method for reducing CHD risk in these people.

## Introduction

Greater body mass index (BMI) is a risk factor for a wide range of adverse health outcomes, including coronary heart disease (CHD), the leading cause of death worldwide. Whilst preventing overweight and obesity is an important public health aim, the substantial and increasing number of people with a high body mass index highlights the need for secondary prevention that aims to reduce risk of the main disease outcomes of high BMI, such as CHD, in those with high BMI, by targeting causal mediators. This is also important because, beyond bariatric surgery^1^ there are no effective and sustainable treatments for those who are obese.

Large prospective population studies show that higher BMI is associated with adverse blood lipid levels, higher fasting glucose, insulin, type 2 diabetes mellitus (T2DM), and CHD. Randomized controlled trials (RCTs) show that elevated triglycerides, low-density lipoprotein cholesterol (LDL-C), glucose and blood pressure increase the risk of CHD.^2,3^ Thus, the association of BMI with CHD could be mediated by these established modifiable risk factors. However, the common method used to test for mediation by observing how much the confounder adjusted multivariable association between a risk factor (e.g. BMI) and outcome (e.g. CHD) reduces with further adjustment for potential mediators,^4^ has been shown to be biased in many situations.^5^

Mendelian randomization (MR), the use of genetic variants as instrumental variables to test the causal effect of risk factors on outcomes, is unlikely to be biased by the extensive confounders of multivariable observational analyses, is less prone to measurement error, and because genetic variants are fixed at conception cannot be biased by reverse causality.^6,7^ As such, it has been used increasingly over the past decade to provide more robust estimates for the causal effect of many risk factors on a range of health outcomes, with results from MR closely resembling those from RCTs where both are available (e.g. the effect of LDL-C^8^ and systolic blood pressure^9^ on CHD). More recent methods have been developed for its use in testing causal mediation using a two-step approach that are considerably less prone to the biases inherent in the common multivariable approach.^5^ Box 1 provides a brief description of MR and its assumptions.

### Box 1: Summary of Mendelian randomisation and its assumptions

Figure Box 1 below shows the assumptions underlying Mendelian randomization

1. *Figure Box 1:Mendelian randomization*

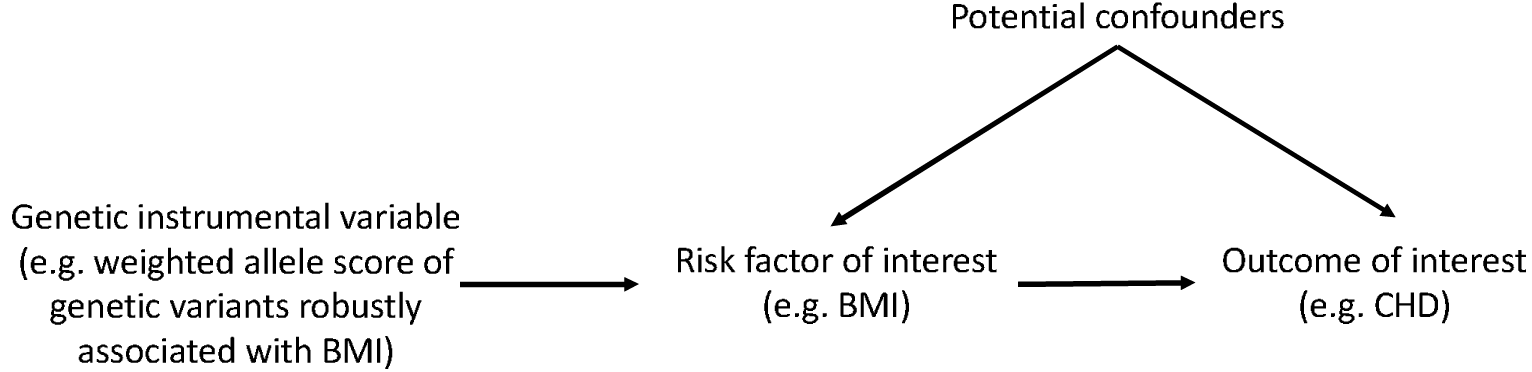 These are that:

- The genetic instrumental variable(s) are robustly related to the risk factor of interest (here BMI); this is illustrated in the Figure above by the arrow from the genetic instruments to BMI
- There is no relationship between any confounders of the risk factor (BMI) and outcome (CHD) and the genetic instrumental variable; illustrated by the lack of any arrow between these confounders and the genetic instrument
- There is no path from the genetic instrument to the outcome, other than through its relationship to the risk factor; illustrated by the lack of any arrow that goes directly from the genetic instrument to the outcome. Empirical evidence suggests that the most likely of these three assumptions to be violated, and result in potentially biased results, is the last one. This may be violated in MR studies by horizontal pleiotropy– that is where the genetic instrument(s) affect other factors which independently of their impact on the risk factor of interest, influence the outcome. If this horizontal pleiotropy is present then the MR estimate of the effect of a risk factor on outcome will be biased, it will actually be the combined effect of that risk factor and any other (pleiotropic) paths from the genetic instruments to outcome. The bias could be an exaggeration of the true effect (if the horizontal pleiotropic paths are in the same direction as that of the main risk factor of interest) or a diminution of the true effect (if the horizontal pleiotropic effect is in the opposite direction of the risk factor of interest).
2. *Estimating causal effects in MR* There are a number of different statistical methods that can be used to estimate causal MR effects. Many of these are related to the ratio, which is intuitive. If the assumptions above are correct then the causal effect of the risk factor (BMI) on outcome (CHD) is the ratio of ‘the association of genetic instruments with CHD’ to ‘the association of genetic instrument with BMI’.
3. *Two sample MR* Valid MR estimates can be obtained using two (independent) samples for the association of the genetic instrument with outcome and the association of genetic instrument with risk factor.^15^ There are some advantages of this 2-sample MR approach over the 1-sample approach (where both parts of the ratio are obtained from the same sample), including the potential to gain very large sample sizes by using publicly available aggregate genome-wide data as we have done here and apply novel methods for testing horizontal pleiotropy that have been developed for use in 2-sample MR with aggregate GWAS data (see methods section of paper and supplementary material for detailed descriptions of these).

Previous MR studies using data from three collections have shown that higher BMI increases risk for CHD (**Supplementary Figure 1** shows results of our meta-analysis of these previous MR studies).^10–13^ These studies used one-sample MR and were unable to undertake sensitivity analyses that have recently been developed for testing likely bias by pleiotropy (Box 1).^14^ The number of cases of CHD varied from 3062 to 11056, which is modest for MR studies. Although MR is likely to be less biased than conventional multivariable approaches, it usually requires a large sample size. Only one of these MR studies analysed potential mediators of the impact of BMI on CHD. It concluded that LDL-C, remnant cholesterol and systolic blood pressure, explained 8%, 7% and 7%, respectively, of the effect of BMI on CHD.^13^ That study was unable to explore potential mediation by insulin sensitivity or hyperglycaemia, which are strongly influenced by BMI and are strong risk factors for CHD. Here, we aimed to investigate the mediating effect of lipid and insulin/glycaemic traits of the effect of BMI on CHD using a large MR study, including over 60,000 CHD, and to analyse a wider set of potential mediators including glycaemic traits (fasting glucose and insulin, Hb_A1c_, T2DM) than previous studies.

## Methods

We used two-step two-sample MR^5,15^ with publicly available datasets that provide genome-wide association results for BMI, glycaemic traits, lipids and CHD. Two-sample MR refers to the use of different datasets (samples) to obtain the gene-risk factor (e.g. BMI) and gene-outcome (e.g. CHD) associations. We firstly tested the effects of BMI on CHD, and then the effects of potential mediation using two-step MR. In step-one we tested causal effects of BMI on potential mediators and in step-two the causal effects of potential mediators on CHD.^5^

### Data sources

#### (a) Genetic instrumental variable for BMI

From the most updated genome wide associations studies (GWAS) on BMI, we obtained 77 single nucleotide polymorphisms (SNPs), identified from the primary meta-analysis of 322,154 European-descent individuals, independently contributing to BMI at genome wide significance (p<5×10^−8^).^16^ These variants were defined as being independent of each other on the basis of low correlation (R^2^ < 0.1) in HapMap22 or the 1000 genome project data. These 77 SNPs account for 2.4% of BMI phenotypic variance.^16^ As a sensitivity analysis we further included 20 SNPs from the secondary analysis of this GWAS;^16^ these include some SNPs that did not reach genome wide significance in Europeans.

#### (b) Potential mediators

Association of SNPs with the phenotypes were extracted from publicly available GWAS consortia. Data on type 2 diabetes mellitus (T2DM) GWAS correlates was obtained from the DIAbetes Genetics Replication And Meta-analysis (DIAGRAM, http://diagram-consortium.org/downloads.html). which includes 34,840 cases and 114,981 controls of European origin.^17^ Genetic associations with fasting insulin (n=38,238), fasting glucose (n=46,186) and glycosylated haemoglobinA_1c_ (HbA_1c_) (n=46,368) were obtained from the Meta-Analyses of Glucose and Insulin-related traits Consortium (MAGIC) and were downloaded from http://www.magicinvestigators.org/; participants were of European ancestry without diabetes.^18^ Genetic associations with high density lipoprotein cholesterol (HDL-C), LDL-C and triglycerides in 188,577 Europeans were obtained from the Global Lipids Genetics Consortium (GLGC) investigators and were downloaded from http://csg.sph.umich.edu/abecasis/public/lipids2013/.^19^

#### (c) Study outcome: coronary heart disease

Data on coronary artery disease/myocardial infarction were obtained from the Coronary ARtery DIsease Genome wide Replication and Meta-analysis (CARDIoGRAM) plusC4D investigators and have been downloaded from www.CARDIOGRAMPLUSC4D.ORG.^20^ This includes 60,801 CHD cases and 123, 504 controls. We first searched the CARDIoGRAMplusC4D 1000 Genomes-based GWAS, a meta-analysis of GWAS studies of mainly European, South Asian, and East Asian, descent imputed using the 1000 Genomes phase 1 v3 training set with 38 million variants.^21^ If no summary data on the gene-CHD association were found from the 1000 Genomes, we screened in CARDIoGRAMplusC4D Metabochip next. If the targeted SNPs were not found in either the 1000 Genomes and the CARDIoGRAMplusC4D Metabochip, we then screened CARDIoGRAM GWAS. The genetic variants used as instrumental variables for CHD, BMI and CHD risk factors (potential mediators) are all shown in **Supplementary Tables 1 to 9.**

### Statistical analysis

As an indication of the strength of the association between genetic instruments and phenotypes, we report the proportion of variation in BMI and all mediators explained by their genetic variant instruments and also the F-statistic for the regression of BMI and all mediators on their genetic instruments.

The proportion of the BMI-CHD effect that is explained by a group of mediators will be estimated with bias if the mediators are related to each other and also if the outcome has an effect on the mediator (i.e. there is reverse causality). Therefore, we tested for potential bi-directional causal effects of BMI, potential mediators and CHD with each other using the inverse variance weighted (IVW) approach described below.

To provide comprehensive assessment using two-sample instrumental variable analysis, five different analytical approaches were used for both step-one (effect of BMI on CHD and potential mediators) and step-two (effect of potential mediators on CHD) of the two-step MR mediation approach. Each of the five methods are different approaches that can be used in the two-sample MR framework. The value of comparing results from all five is that they have some different underlying assumptions and therefore we have more confidence in results that are consistent across the different methods. Full details of these approaches, including their different assumptions, are provided in the Supplementary Methods.

To estimate the effect of BMI on CHD taking account of genetically determined potential mediators, we used the IVW MR method, adjusting for the SNP-potential mediator effect.^22^ The proportion of the effect that is mediated by any of the potential mediators was estimated by the changes in the total effect of the genetically determined BMI on CHD risk (more details in the Supplementary Methods). This method assumes that mediators are continuously measured variables and as T2DM is dichotomized we did not assess the proportion of the BMI-CHD effect due to T2DM.

A detailed analysis diagram was shown in Figure 1. All statistical analysis was performed using STATA 13.1 and R-software (Version 3.2.5).

**Figure 1.**
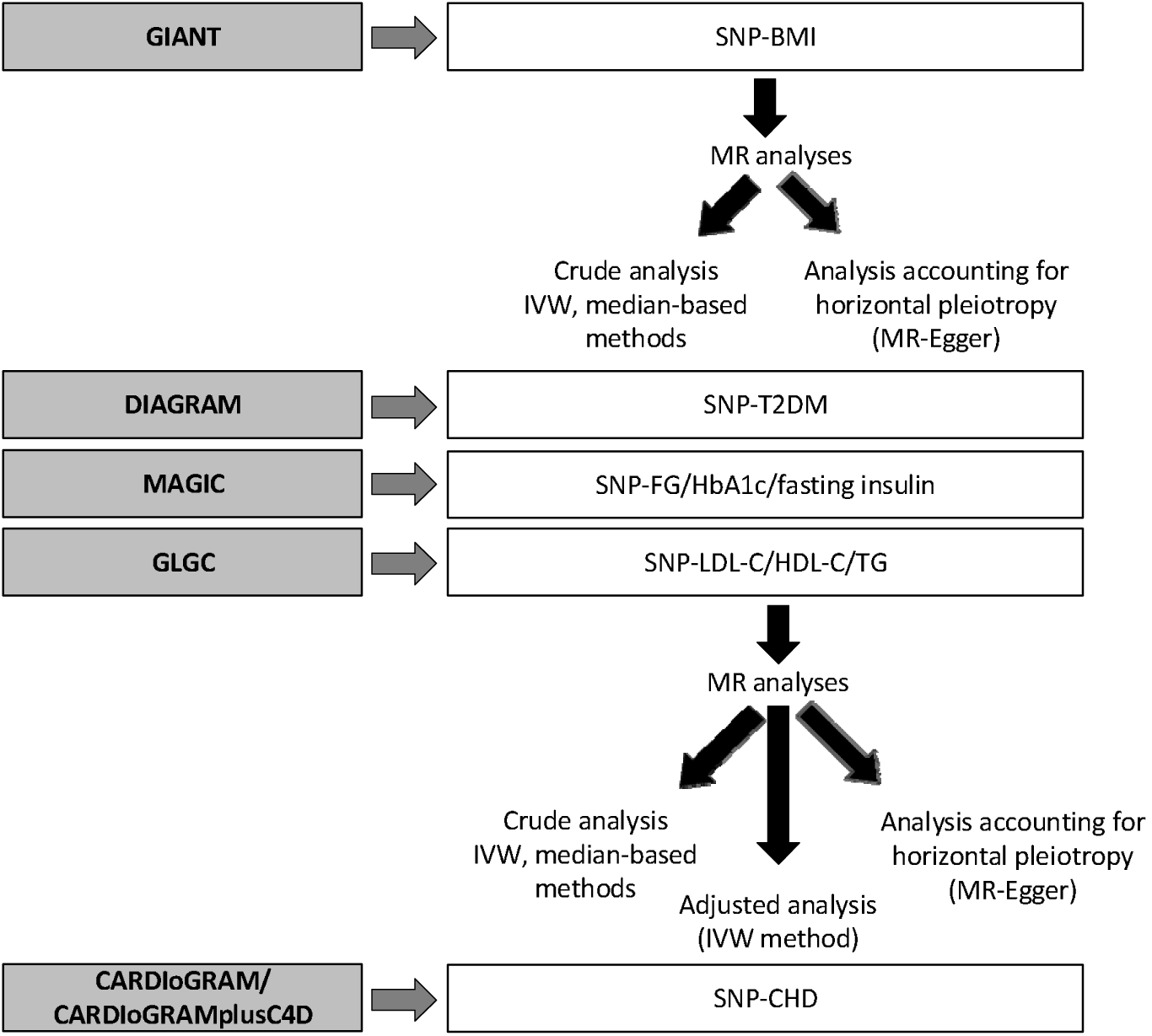
Analysis diagram. Summary data for SNP-phenotypes were extracted from GWAS consortia datasets (GIANT, CARDIoGRAM, C4D, DIAGRAM, MAGIC and GLGC). MR estimates of BMI on mediators (T2DM, fasting glucose, fasting insulin, HbA_1c_, LDL-C, HDL-C and triglycerides), and of BMI and mediators on CHD were derived using IVW method. SNPs: single nucleotide polymorphisms; GIANT: Genetic Investigation of Anthropometric Traits; CARDIoGRAM: Coronary Artery Disease Genome-wide Replication and Meta-analysis;CARDIoGRAMplusC4D Metabochip: CARDIoGRAMplusC4D Metabochip meta-analysis; DIAGRAM: DIAbetes Genetics Replication And Meta-analysis; MAGIC: Meta-Analyses of Glucose and Insulin related traits Consortium; GLGC: Global Lipids Genetics Consortium; BMI: body mass index; HbA_1c_: glycated hemoglobin; LDL-C: low density lipoprotein cholesterol; HDL: high density lipoprotein cholesterol; CHD: coronary heart disease; IVW method: inverse variance weighted method; MR: Mendelian randomization.

## Results

The proportion of variation explained by all of the variants that we used as instrumental variables for the potential mediators varied from 1.2% (for fasting insulin) to 5.7% (for T2DM) (Supplementary Tables 1-8). The first stage F-statistic for all of the MR analyses (i.e. for the regression of BMI and each of the mediators on their genetic variant instrument variables) were all very large (> 500).

### Relationships between potential mediators and CHD

As expected, we observed evidence for bidirectional association between FPG and T2DM, and for both FPG and T2DM associate with HbA_1c_ (Table 1). LDL-C, HDL-C and triglycerides appeared related to each other. CHD appears to be causally positively related to T2DM, but was not related to other potential mediators (Table 1).

**Table 1.** Mendelian randomization estimates^†^ of risk factors with each other and with coronary heart disease and type 2 diabetes

| Exposures | BMI, SD | T2DM | FPG, mmol/l | HbA<sub>1c</sub>, % | FI, log-pmol/l | LDL-C, SD | HDL-C, SD | TG, SD | CHD |
| --- | --- | --- | --- | --- | --- | --- | --- | --- | --- |
| **Outcomes** |  |  |  |  |  |  |  |  |  |
| BMI, SD | 0 | -0.07 (0.009)*** | 0.007 (0.04) | 0.04 (0.05) | -0.51 (0.14)*** | -0.04 (0.01)* | 0.0008 (0.01) | -0.004 (0.02) | -0.015 (0.01) |
| T2DM | 0.67 (0.19)*** | 0 | 1.23 (0.14)*** | 0.36 (0.2) | 0.53 (0.45) | -0.09 (0.06) | -0.03 (0.05) | 0.05 (0.08) | 0.10 (0.05)* |
| FPG, mmol/l | 0.07 (0.02)*** | 0.08 (0.007)*** | 0 | 0.07 (0.07) | 0.05 (0.07) | -0.04 (0.01)** | -0.002 (0.01) | -0.01 (0.02) | 0.01 (0.01) |
| HbA<sub>1c</sub>, % | 0.05 (0.03)*** | 0.03 (0.01)** | 0.45 (0.04)*** | 0 | -0.09 (0.1) | -0.01 (0.02) | -0.02 (0.02) | -0.03 (0.02) | 0.01 (0.02) |
| FI, log-pmol/l | 0.18 (0.03)*** | 0.06 (0.009)*** | 0.05 (0.05) | -0.06 (0.06) | 0 | -0.03 (0.02) | 0.007 (0.02) | 0.02 (0.02) | -0.008 (0.01) |
| LDL-C, SD | -0.05 (0.07) | 0.02 (0.01) | 0.02 (0.04) | 0.09 (0.05) | 0.27 (0.14)* | 0 | -0.21 (0.02)*** | 0.19 (0.03)*** | -0.03 (0.02) |
| HDL-C, SD | -0.23 (0.04)*** | -0.003 (0.01) | 0.04 (0.04) | 0.11 (0.05)* | 0.52 (0.16)*** | -0.18 (0.02)*** | 0 | -0.47 (0.03)*** | -0.02 (0.02) |
| TG, SD | 0.20 (0.03)*** | 0.02 (0.01) | 0.03 (0.03) | -0.02 (0.05) | -0.21 (0.16) | 0.07 (0.03)* | -0.16 (0.01)*** | 0 | 0.001 (0.02) |
| CHD | 0.37 (0.07)*** | 0.10 (0.03)*** | 0.19 (0.09)* | 0.31 (0.12)* | -0.49 (0.31) | 0.49 (0.05)*** | -0.13 (0.04)** | 0.21 (0.05)*** | 0 |

BMI: body mass index; FPG: fasting plasma glucose; LDL-C: low-density lipoprotein cholesterol; HDL-C: high-density lipoprotein cholesterol; TG: triglycerides; FI: fasting insulin; CHD: coronary heart disease; HbA<sub>1c</sub>: glycosylated haemoglobin A<sub>1c</sub>; T2DM: type 2 diabetes mellitus
<sup>†</sup>: All results are beta coefficients (standard error) from the MR instrumental variable estimates using inverse variance weighted and so reflect differences in mean outcome per 1 unit difference of the exposures for continuously measured outcomes and difference in log odds for binary outcomes (CHD/T2DM)
\* : P<0.05; \*\* : P<0.01; \*\*\* : P<0.001

### Effects of BMI on CHD and glycaemic and lipid traits

The IVW MR method showed evidence that BMI was causally related to CHD and all of the glycaemic and lipid traits except LDL-C (Table 2). There was consistent support across all five MR methods for a causal effect of greater BMI on increased CHD and T2DM risk and levels of fasting glucose, HbA_1c_, fasting insulin and triglycerides, together with decreased HDL-C. None of the methods supported a causal effect of BMI on LDL-C (Table 2 **and Supplementary Table 10**).

**Table 2.**
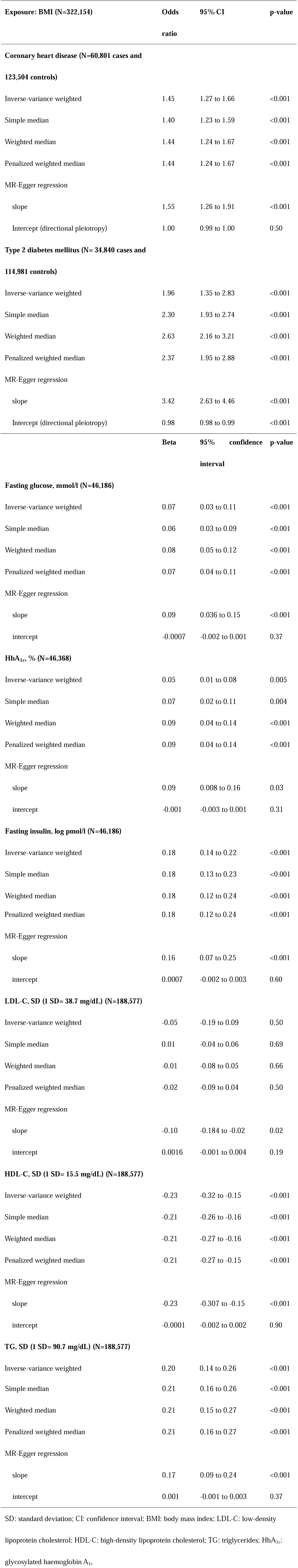
Mendelian randomization estimates of body mass index (SD, 1 SD=4.5 kg/m^2^) on cardiovascular risk factors and coronary heart disease

### Effects of potential mediators on CHD

There was broadly consistent support across all five MR methods for a positive effect of T2DM, HbA_1c_, triglycerides and LDL-C on CHD risk (Table 3). For T2DM the MR-Egger 95% confidence interval just included the null value, but this method has lower statistical power than the other four methods and the point estimates were similar for all five methods. For triglycerides the estimate of effect (slope) from MR-Egger was a little weaker than for all of the other methods (e.g. 1.24 versus 1.13 comparing the IVW and MR-Egger methods), suggesting that some, but not all, of the effect of triglycerides estimated by IVW and other methods might be due to horizontal pleiotropy. In IVW, and the median methods analyses, lower HDL-C and higher fasting glucose and insulin appeared to be causally related to higher risk of CHD. However, for all of these MR-Egger suggested that effects were largely due to horizontal pleiotropy, with effect estimates markedly attenuated to the null and the intercepts all being non-zero.

**Table 3.**
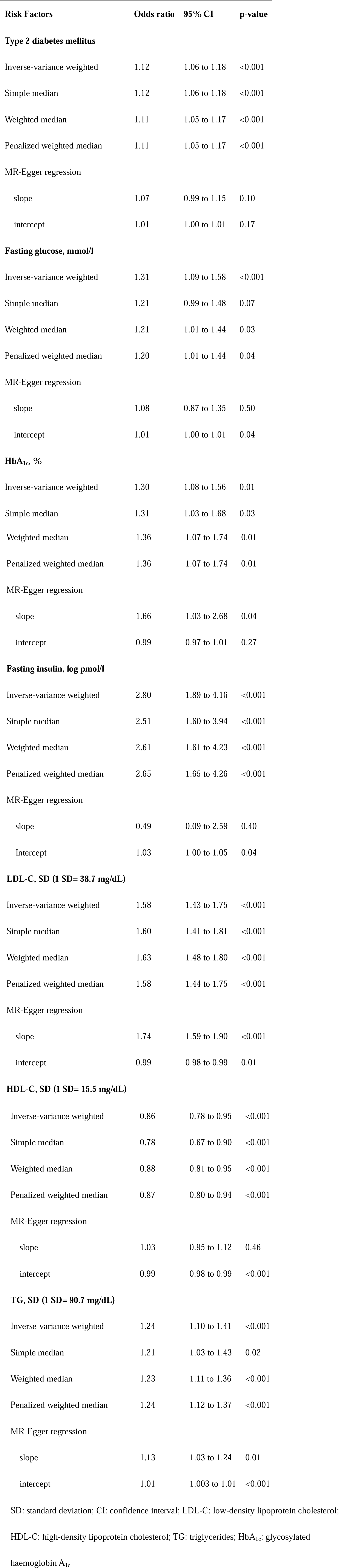
Mendelian randomization estimates of cardiovascular risk factors on coronary heartdisease

### Mediating effects of lipids and glycaemic traits on BMI-CHD effects

We explored those potential mediators that had causal MR support for both an effect of BMI on them (step-one) and of the mediators on CHD (step-two): T2DM, HbA_1c_ and triglycerides. Our results suggested that triglycerides were an important mediator, with either T2DM or HbA_1c_ further contributing to mediation of BMI on CHD. The BMI-CHD effect reduced from 1.45 (95% confidence interval (CI): 1.27, 1.66) to 1.16 (95% CI: 0.99, 1.36) and to 1.36 (95% CI: 1.19, 1.57) with genetic adjustment for triglycerides or HbA_1c_ respectively, and to 1.09 (95% CI: 0.94, 1.27) with adjustment for both (Table 4 **and Supplementary Table 11**).

**Table 4.** Multivariate separate-sample Mendelian randomization analysis of the effect of body mass index (per SD, 1 SD=4.5 kg/m^2^) on coronary heart disease

|  | **Odds ratio** | **95% CI** | **p-value** | **Mediation effect** |
| --- | --- | --- | --- | --- |
| MR-IVW regression, crude | 1.45 | 1.27 to 1.66 | <0.001 |  |
| Multivariate model |  |  |  |  |
| (1) adjusted for TG | 1.16 | 0.99 to 1.36 | 0.06 | 22% |
| (2) adjusted for HbA<sub>1c</sub> | 1.36 | 1.19 to 1.57 | 0.001 | 4% |
| (3) adjusted for T2DM | 1.35 | 1.17 to 1.56 | 0.001 | - |
| (4) adjusted for TG+HbA<sub>1c</sub> | 1.09 | 0.94 to 1.27 | 0.25 | 38% |
| (5) adjusted for TG+ T2DM | 1.10 | 0.94 to 1.29 | 0.22 | - |

SD: standard deviation; CI: confidence interval; TG: triglycerides; HbA<sub>1c</sub>: glycosylated
haemoglobin A<sub>1c</sub>; T2DM: type 2 diabetes mellitus

## Discussion

We used MR analysis to investigate the extent to which glycaemic and/or lipid traits might mediate a causal path between BMI and CHD. Consistent with previous studies,^10–13^ but here using a larger number of CHD cases and genetic variants, we show that higher BMI causes greater CHD risk. Our results also suggest that triglycerides, HbA_1c_ and T2DM play important roles in causally mediating the effect of BMI on CHD. By contrast our results suggest that BMI is not causally related to LDL-C and that HDL-C, fasting glucose and insulin may not be causally related to CHD. Thus, our findings provide strong support for undertaking RCTs in obese people to test the effect of triglyceride reduction and glycaemic control on CHD risk.

Several MR studies have previously examined the association of BMI with CHD and CHD risk factors.^10,11,13,23,24^ Our results are broadly consistent with those, including our finding that BMI does not appear to be causally related to LDL-C.^11,23,25^ This is further supported by two RCTs of bariatric surgery which find intensive weight loss does not lower LDL-C.^26,27^ Consistent with our results, previous MR studies have also shown positive causal effects of T2DM, HbA_1c_, LDL-C and triglycerides, with CHD^28,29,30,31^, but not fasting glucose or HDL-C once horizontal pleiotropy has been accounted for. The discrepancy between finding a causal effect of T2DM and HbA_1c_ on CHD, but not of fasting glucose, might suggest that non-fasting (post-prandial) glucose, more so than fasting levels, are most relevant for CHD risk, and/or that long-term hyperglycaemia (as assessed by elevated HbA_1c_ and likely to be identified as being above the threshold required to diagnose T2DM) are important.

To our knowledge only one previous study has tried to explore potential mediation of the BMI-CHD effect in an MR framework. That study included 11,056 CHD cases and 75,627 controls from Copenhagen and used only 3 BMI-related SNPs. It concluded that the effect of BMI on increased CHD risk was partly mediated through elevated levels of LDL-C, non-fasting remnant cholesterol and systolic blood pressure.^13^ The evidence for a mediating role of remnant cholesterol is entirely consistent with our findings here for triglycerides, as remnant cholesterol is the cholesterol content of triglyceride-rich lipoproteins, particularly so in this previous study where remnant cholesterol was not directly measured but estimated from other lipids using a method that would produce an extremely high correlation between (measured) triglycerides and estimated remnant cholesterol.^13^

None of our study, the previous (Copenhagen) study,^12,13^ other MR studies^11,23,25^ or RCTs of bariatric surgery^32,33^ have found strong evidence for a causal effect of BMI on LDL-C, which suggests it is unlikely to be an important mediator of BMI on CHD. However, since the previous study (despite finding no MR evidence for a causal effect of BMI on LDL-C) concluded that LDL-C was a partial mediator^13^ we examined that possibility in our data. As expected we found no strong support for a mediating effect of LDL-C between BMI and CHD (**Supplementary Table 12**). We were unable to explore any mediating effect of blood pressure in our study. This is because our approach uses publicly available aggregate genome-wide results and the International Consortium for Blood Pressure (ICBP) provides information on SNPs and blood pressure associations without specifying the risk (or effect) allele for each SNP, and thus the effect of BMI on blood pressure cannot be assessed using two-sample MR instrumental variable analysis.

CHD is a major cause of morbidity and mortality and its prevalence is increasing worldwide, partly because of the increasing prevalence of obesity. Our results indicate the extent to which acting on risk factors, such as triglycerides, HbA_1c_ and T2DM, might counteract the detrimental effects of obesity on CHD. They highlight the potential importance of using interventions that lower triglycerides and/or HbA_1c_ and T2DM specifically in those with obesity.^34,35^ There is evidence, including from MR, that statins affect triglycerides and remnant cholesterol, as well as LDL-C.^36^ Furthermore, a rare variant in *APOC3* with a marked effect on triglyceride levels provides a potential target for drug development aimed at reducing triglyceride levels, independently of any statin effects.^37,38^ Thus, targets for reducing triglycerides exist and testing the effect of these in obese populations would be feasible. Previous RCTs have shown that the oral hypoglycaemic metformin reduces cardiovascular risk factors^39–41^ in non-diabetic at risk populations, including those who are obese, but its effect on CHD risk has yet to be established. Our results suggest that it might be cardio-protective in populations with high BMI and supports RCTs to test its effect on CHD in these people.

### Strengths and limitations

Our study is extremely large and uses genetic variants to avoid some of the key limitations of traditional multivariable regression approaches to mediation. Horizontal pleiotropy is one of the major concerns in relation to limitations of MR studies. However, to explore the potential effects of this pleiotropy, we used different MR methods (IVW, median-based estimators and MR-Egger) that have different assumptions and we assessed the consistency across each of these estimators. The mediators that we took forward into MR-based mediation analyses (triglycerides, HbA_1c_ and T2DM) had consistent causal effects across these different methods for both steps – i.e. the effect of BMI on them and of them on CHD. In the mediation analyses, where we include both genetically predicted triglycerides and HbA_1c_, we are assuming that these two (triglycerides and HbA_1c_) are not causally related to each other. We tested for causal relationships between potential mediators prior to our main two-step MR analyses and these suggested that triglycerides are not causally related to HbA_1c_ or other glycaemic traits. However, MR studies cannot completely rule out a causal relationship between the two. Previous large prospective studies showed triglycerides predicted the development of T2DM,^42,43^ if this association is casual, the estimated mediation effect by dysglycaemia and triglycerides could be inflated. Our results would be biased if the mediators we have tested caused variation in BMI (i.e. there was reverse causality from mediators to BMI). If this were the case, we would expect a bi-directional MR effect between BMI and the mediating risk factors. However, we found no evidence that triglycerides or HbA_1c_ caused variation in BMI (though the causal effect of BMI to these mediators was present).

Whilst all five MR methods suggest a casual effect of triglycerides on CHD, the MR-Egger intercept suggest that there might be some horizontal pleiotropy for this effect. It is plausible that the genetic variants we used as instruments for triglycerides also affect other remnant cholesterols or other lipids and in fact these also contribute to mediating BMI effects on CHD. Another potential limitation to our study is that we have assumed no interaction between BMI and mediators, but we are not able to test for this because we have used aggregated genome-wide data. Previous observational studies suggest that the association between BMI and CHD may be modified by hypertension,^44^ but did not find effect modification by the glycaemic and lipid traits that we have examined here.^45^ In two sample MR weak instrument bias can result in bias towards the null. In mediation analyses this could result in an underestimation of mediating effects. However, given our large sample size and the fact that our genetic instruments explained 2.1% and 2.4% of the variation in triglycerides and HbA_1c_, respectively, and had very large first-stage F-statistics, we think this is unlikely to have had a major effect on our results.

In conclusion, our result support a causal effect of higher BMI increasing risk of CHD, that is, at least partially, mediated through effects of BMI on triglycerides, HbA_1c_ and T2DM. These findings support the need for interventional studies examining whether lowering triglycerides or providing anti-diabetic therapy in people who are overweight or obese is effective at reducing their increased risk (in comparison to healthy weight people) of CHD.

## Acknowledgements

Data on body mass index have been contributed by Genetic Investigation ofANthropometric Traits (GIANT) consortium and have been downloaded from http://www.broadinstitute.org/collaboration/giant/index.php/CIANT_consortium_data_files. Data on coronary artery disease/myocardial infarction have been contributed by CARDIoGRAMplusC4D investigators and have been downloaded from www.CARDIOGRAMPLUSC4D.ORG’. Data on lipid traits have been contributed by Global Lipids Genetics Consortium and have been downloaded from http://csg.sph.umich.edu/abecasis/public/lipids2013/. Data on glycaemic traits have been contributed by MAGIC investigators and have been downloaded from www.magicinvestigators.org. All of these data are publicly available.

Investigators who have made their genome-wide data available to scientist may not necessarily agree with comments made in this paper and the authors take full responsibility for the contents of this paper.

## Author contributions

Study design: DAL, LX, MCB, GH Analysis plan: LX, DAL, MCB Data acquisition (from public data): LX, GH Analyses: LX Writing first draft of paper: LX Critical comments and contributions to final writing of paper: DAL, MCB, GH

